# Contributions of early and mid-level visual cortex to high-level object categorization

**DOI:** 10.1101/2023.05.31.541514

**Authors:** Lily E. Kramer, Yi-Chia Chen, Bria Long, Talia Konkle, Marlene R. Cohen

## Abstract

The complexity of visual features for which neurons are tuned increases from early to late stages of the ventral visual stream. Thus, the standard hypothesis is that high-level functions like object categorization are primarily mediated by higher visual areas because they require more complex image formats that are not evident in early visual processing stages. However, human observers can categorize images as objects or animals or as big or small even when the images preserve only some low- and mid-level features but are rendered unidentifiable (‘texforms’, Long et al., 2018). This observation suggests that even the early visual cortex, in which neurons respond to simple stimulus features, may already encode signals about these more abstract high-level categorical distinctions. We tested this hypothesis by recording from populations of neurons in early and mid-level visual cortical areas while rhesus monkeys viewed texforms and their unaltered source stimuli (simultaneous recordings from areas V1 and V4 in one animal and separate recordings from V1 and V4 in two others). Using recordings from a few dozen neurons, we could decode the real-world size and animacy of both unaltered images and texforms. Furthermore, this neural decoding accuracy across stimuli was related to the ability of human observers to categorize texforms by real-world size and animacy. Our results demonstrate that neuronal populations early in the visual hierarchy contain signals useful for higher-level object perception and suggest that the responses of early visual areas to simple stimulus features display preliminary untangling of higher-level distinctions.

## Introduction

The complexity of the visual features for which neurons are tuned increases along the ventral visual stream (Felleman & Van Essen, 1991; Mishkin et al., 1983). This increase is mimicked in subsequent layers of deep artificial neural networks optimized for object recognition (Khaligh-Razavi & Kriegeskorte, 2014; Yamins et al., 2014; Yamins & DiCarlo, 2016). As such, it has been regularly hypothesized that high-level functions like object categorization are primarily mediated by later areas in the visual hierarchy (Biederman, 1987; DiCarlo & Cox, 2007; Krizhevsky et al., 2012; Riesenhuber & Poggio, 1999). Specifically, the prevailing hypothesis holds that because neurons in earlier visual areas encode simple visual features, the representations of, for example, images of animals are ‘entangled’ with those of objects in earlier but not later stages of visual processing. The assumption of this hypothesis is that images of animals and images of objects do not have enough distinguishing featural information to be linearly separable at early visual encoding stages, and only become separable later in the ventral stream, requiring more hierarchical featural processing (DiCarlo & Cox, 2007; Grill-Spector & Weiner, 2014; Ungerleider & Bell, 2011).

Despite the popularity of the late-stage hypothesis, an emerging body of evidence suggests that low- or mid-level features already carry enough information regarding animacy and real-world size to directly influence human behaviors (Grootswagers et al., 2019; S. P. Li & Bonner, 2021; Lieber et al., 2023; Long et al., 2017; Wang et al., 2022). For example, people are able to distinguish animals from inanimate objects, and big from small items (in an identical viewing size) above chance performance, when only some low- and mid-level visual features are available (Long et al., 2016, 2017, 2018). To isolate the low- and mid-level visual features, these studies invented and used ‘texform’ stimuli. Texforms are unrecognizable images created using a spatially-constrained texture synthesis model (Freeman & Simoncelli, 2011; Oleskiw et al., 2020) which preserves correlations between mid-level image statistics across different spatial scales (Long et al., 2017). Additionally, recent studies have also shown that humans’ and monkeys’ ability to judge animacy is linked to the amount of image-based curvilinearity information contained in the stimuli (Zachariou et al., 2018; Yetter et al., 2021).

Multiple physiological studies have also provided evidence supporting the role of low- and mid-level features in high-level functions, primarily by noting that the patterns of neuronal selectivity to lower-level features might be useful for high-level functions. For example, the differing curvature tuning for neurons with receptive fields at the fovea (preferring curvy features) and in the periphery (preferring boxy features) may support the specialization of face recognition at the fovea and landscapes in the periphery (Arcaro & Livingstone, 2017; Hasson et al., 2002; Kravitz et al., 2013). In studies specific to animacy and real-world size, it was shown that linear decoders were sufficient to make above chance classifications from EEG and fMRI signals evoked by mid-level features from texforms (Long et al., 2018; Wang et al., 2022).

Moreover, there is electrophysiological evidence that information about the animacy of images (with all visual features intact) is accessible early in the population responses of V4 neurons (Cauchoix et al., 2016).

We reasoned that a strong test of the hypothesis that low- and mid-level feature differences are sufficient for the preliminary untangling of representations of image animacy and real-world size was to 1) measure information related to object categorization in populations of neurons in early and mid-level visual cortex in monkeys and 2) relate that activity to perception. Therefore, we recorded the responses of a few dozen neurons in primary visual cortex (V1) and mid-level visual area V4 while rhesus monkeys viewed unaltered images of animals and objects and texforms made from those same images. We then assessed how well we could linearly decode category information from those populations and the extent to which the decoded information relates to human perception.

Even though we recorded from only a small proportion of the thousands of cells in each area that respond to these images, we could readily decode animacy and real-world size from the population responses. Critically, our ability to classify each texform from the responses of neurons in monkey visual cortex was also related to the ability of humans to categorize those same texforms. These results suggest that the low- and mid-level visual feature information encoded by V1 and V4 neurons is sufficient to begin the process of untangling these high-level visual distinctions of animacy and real-world size. Moreover, the relationship between neural decoding and human categorization performance is consistent with the possibility that early and mid-level visual cortical areas play an important role in high-level functions like object classification.

## Results

### Real-world size and animacy information is evident in early and mid-level visual cortex

We presented grayscale images of objects and animals that were either unaltered (Figure 1a, left) or had been generated via a texture synthesis algorithm to preserve only some low- and mid-level visual features (‘texforms’, Figure 1a, right) to three adult male rhesus monkeys (*Macaca mulatta,* weights 10, 10, and 11 kg). While animals viewed these images, we recorded from neural populations in primary visual cortex (V1) or visual area V4 (V1 recordings in monkeys M1 and M3, V4 recordings in monkeys M1 and M3; Figure 1b). We recorded from either chronically implanted multielectrode arrays (monkeys M1 and M2) or a movable 32-channel probe (monkey M3; see Methods). We presented one unaltered image or texform within the joint receptive fields of the recorded units on each trial (Figure 1b). To reduce adaptation between trials, after the image turned off, the monkey was rewarded for making an eye movement to a subsequently presented target.

**Figure 1.**
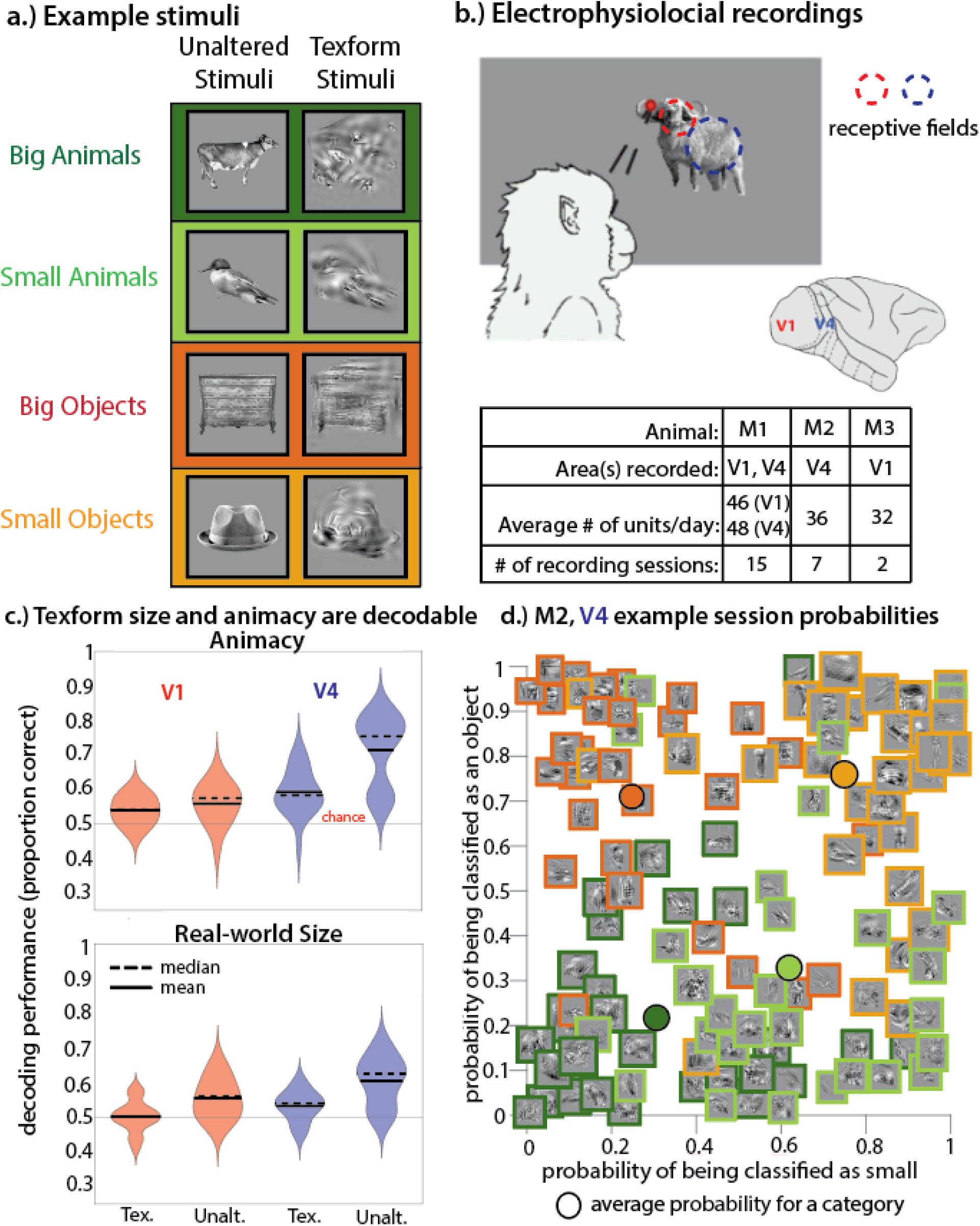
Animacy and real-world size decoding in V1 and V4. **(a) Example stimuli.** A total of 480 stimuli were utilized. Half of those stimuli were the unaltered images (left column) that the texform stimuli (right column) were generated from via a texture synthesis algorithm, as tested in Long et al,. 2018 and Deza et al., 2019. Texform stimuli preserve some low- and mid-level visual features of their unaltered stimuli counterparts within spatially defined pooling windows, but are unrecognizable to human observers. See methods for further explanation. **(b) Electrophysiological recordings.** Stimuli were displayed on a computer screen overlapping the receptive field locations of units in the visual cortical areas recorded (V1 and V4). Table detailing data obtained from each animal (M1, M2, and M3). During a trial, animals were required to fixate for 100-200 ms before a stimulus was presented. Animals maintained fixation during the stimulus presentation period (860 ms). The fixation spot and stimulus then disappeared and a target appeared at a random radial location not overlapping the previous location of the stimulus. Animals were then required to make a saccade to the location of the target in 150-250 ms. **(c) Decoding real-world size and animacy.** Decoding performances for each brain area (V1 in red, V4 in blue) averaged over all sessions (both animals for each area included in mean and median). Width of the violin plot represents the kernel density estimation. All distributions of session decoding performances were significantly above chance performance (one-sided Wilcoxon signed rank tests, p<0.01), except for the condition of decoding real-world size of texforms from V1 population responses. **(d) Example session texform classification probabilities.** The probability that the linear decoder trained and tested on neural responses to texforms classifies each stimulus as small or as an object. Data from one example session from M2, V4. Colored outlines represent the actual categories for each stimulus. Color coded circles represent the average classification probability for each category.

We tested the hypothesis that signals related to animacy and size categorization are present in early and mid-level visual areas, in both unaltered images and texform counterparts. We used cross-validated, linear methods to decode the animacy (Figure 1c, top violin plot) and real-world size (Figure 1c, bottom violin plot) of each stimulus from each group of simultaneously recorded cells in each brain area.

We could robustly decode animacy and real-world size from V4 population activity (note the separability of the four colored categories for an example V4 recording session in Figure 1d) for both stimulus classes (one-sided Wilcoxon signed rank tests, *p<0.01* for animacy and size session decoding performance distributions, Figure 1c). Animacy information for both stimulus classes was also decodable from V1 population activity in numerous sessions (one-sided Wilcoxon signed rank tests, *p<0.01* for V1 unaltered and texform animacy distributions, Figure 1c). Real-world size information of unaltered images was consistently decodable from V1 population activity (one-sided Wilcoxon signed rank test, *p=0.0017* for the unaltered size distribution), however, size information of texforms was not (*p=0.406*, for the texform size distribution, Fig. 1c). Although decoding performance varied across recording sessions (in part because of differences in the number of recorded units, evoked firing rates, stimuli used, and the modest number of image presentations), performances were significantly above chance in the majority of sessions, (except in the case of decoding the size of texform stimuli from V1 activity, for which only 7/17 V1 recording sessions were significant).

In general, mean decoding performance was slightly better for unaltered images than texforms in both V1 and V4 (paired, one-sided t-tests indicated better decoding for unaltered images than texforms, *p<0.01*), except in the case of animacy decoding from V1 population activity (*p=0.149*). The ability to decode animacy and size information from population responses to unaltered images demonstrates that linearly decodable information about real-world size and animacy distinctions is already present in the early and middle parts of the ventral visual stream. The texform results further triangulate the complexity of the feature tuning underlying this information, and support the idea that this separability is not simply due to feedback from downstream object recognition mechanisms reliant on high-level visual features.

### Human observers can classify the size and animacy of texforms

A subset (120 texforms, generated via the accelerated method) of the stimuli utilized to collect electrophysiological data in this experiment were the same as those shown to human participants in previous studies (Deza et al., 2019). In that previous work, human observers were able to classify texforms as animals or objects (animacy, y-axis in Figure 2a) or as having big or small real-world size (size, x-axis in Figure 2a) significantly above chance performance (N = 64 human subjects, average animacy classification score = 0.396, t-test, *p<0.001*; average size classification score= 0.167, t-test, *p<0.001*, chance performance score is 0). On an image by image basis, the performance of human subjects classifying animacy was unrelated to performance classifying size (*r* = 0.19, paired sample t-test, *p>0.05*).

**Figure 2.**
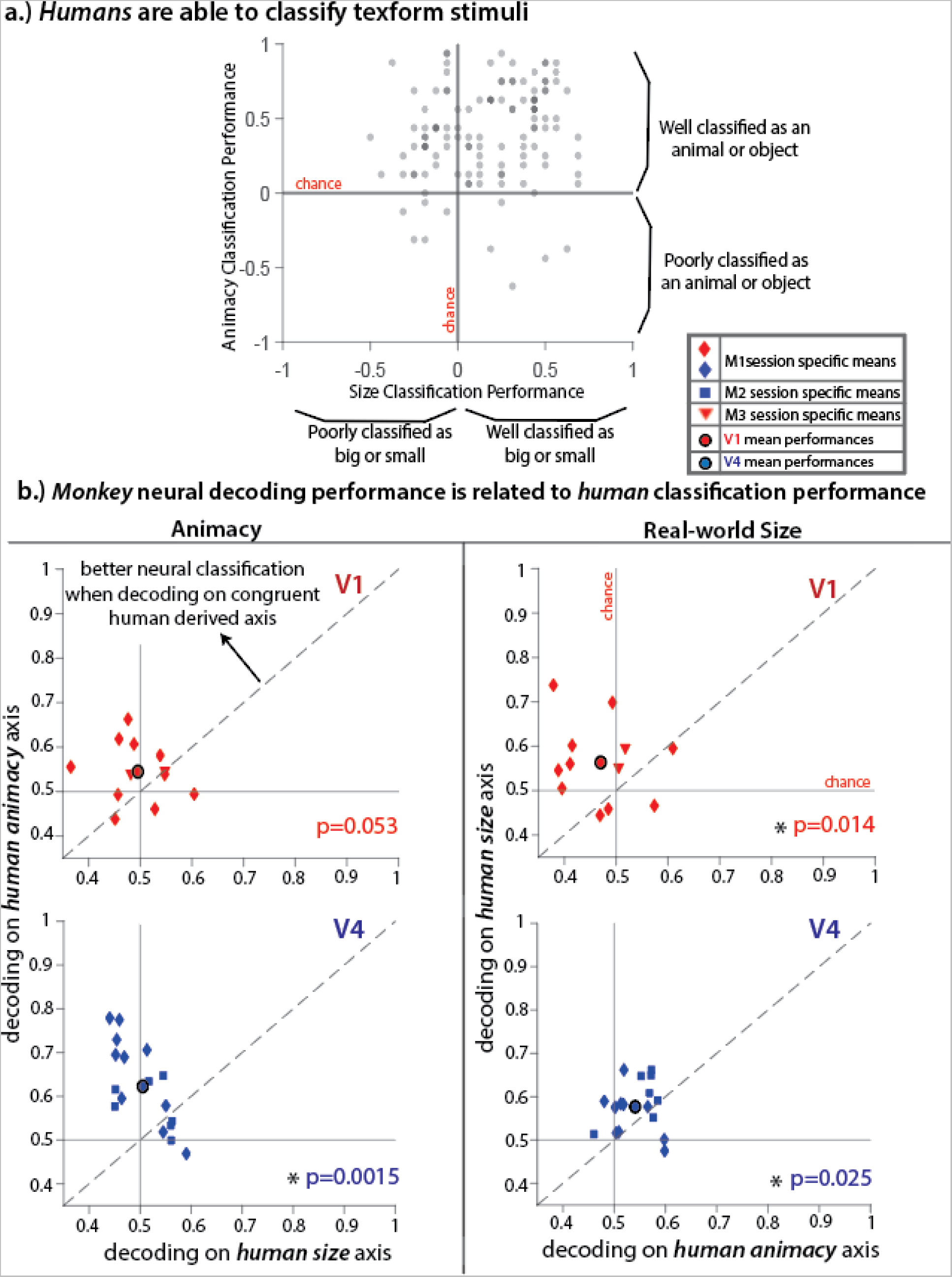
**(a) Human classification scores.** Human classifiability scores for each texform stimulus were previously obtained via online experiments (Chen, Deza, Long, & Konkle., unpublished data). Each point represents one texform stimulus. The higher the score, the more frequently the texform was correctly classified. See methods for further explanation of score calculation. **(b) Monkey neural decoding performance is related to human classification performance.** Mean decoding performances for each recording session (blue and red animal specific symbols) were determined by performing ROC analysis on axes in neural space that best explained either animacy or real-world size human classification scores. Colored circles represent mean decoding performance for each brain area (average across all sessions, V1 red, V4 blue). The y-axes represent decoding performance calculated using the ability of humans to classify the category that was **congruent** to the neural classification dimension (animacy on left, real-world size on right). The x-axes represent decoding performance calculated using the ability of humans to classify the category that was **incongruent** to the neural classification dimension. Asterisks represent conditions where distributions of mean decoding performance for each session are significantly different in the congruent and incongruent scenarios.

### There is a link between neural coding in monkey visual cortex and human perception

To further test the hypothesis that animacy and size categorization could plausibly depend on the low- and mid-level image features encoded by populations of V1 and V4 neurons, we compared the performance of humans categorizing texforms with the performance of our decoder that used linear methods to glean category information from the responses of simultaneously recorded units from monkey V1 or V4. We reasoned that there should be a link between those measures only if categorization behavior and neural responses both reflect the information about the low- or mid-level image features preserved in the texform manipulation. Otherwise, there is no reason to think that human behavior should be related to the responses of a few dozen neurons in a monkey’s brain.

For each analyzed recording session, brain area, and category type (animacy or real-world size), we projected neural population responses onto the axes in neural population space that best explained the ability of human observers to classify texforms according to that category (see Methods for extensive explanation). We compared our ability to decode animacy from neural responses projected onto the human animacy classification axis (congruent condition) with performance decoding responses projected onto the human size axis (incongruent condition), and vice versa for real world size (see Supplemental Figure 2 for schematic illustrating this analysis). If there is a relationship between neural population responses and human classification performance, decoding performance should be better for the congruent conditions (y-axes, figure 2b) than the incongruent conditions (x-axes, figure 2b).

Indeed, we observed a consistent relationship between monkey neural responses and human perception. Our ability to decode animacy or real-world size from monkey neuronal population responses was better along the congruent than the incongruent axes (most points lie above the diagonal line, figure 2b). Decoding performance for the congruent conditions was significantly greater than for the incongruent conditions in nearly all circumstances (one-sided, paired-sample tests, *p<0.05*). Further, for all brain areas, decoding performance for congruent conditions was significantly better than shuffled controls (one-sided, paired-sample t-tests, *p<0.05*; see Methods for details of the shuffle control). These results suggest that high-level object categorization by humans and neurons depend on features of the visual stimuli that are preserved by the texform manipulation.

### Neurons in early and mid-level visual areas respond similarly to unaltered images and their texform counterparts

There was a strong neuron-by-neuron correlation between responses to an unaltered image and its texform counterpart (Figure 3a; *p<0.05* for each monkey, brain area, and session). The activation profile across units in response to an unaltered image was highly correlated with the response to its texform counterpart (red/blue distributions), relative to control comparisons relating the responses between pairs of unaltered images (dark gray distribution) or pairs of texform images (light gray distribution). Thus, the populations of V1 and V4 neurons responded similarly to unaltered images and their texform counterparts.

**Figure 3.**
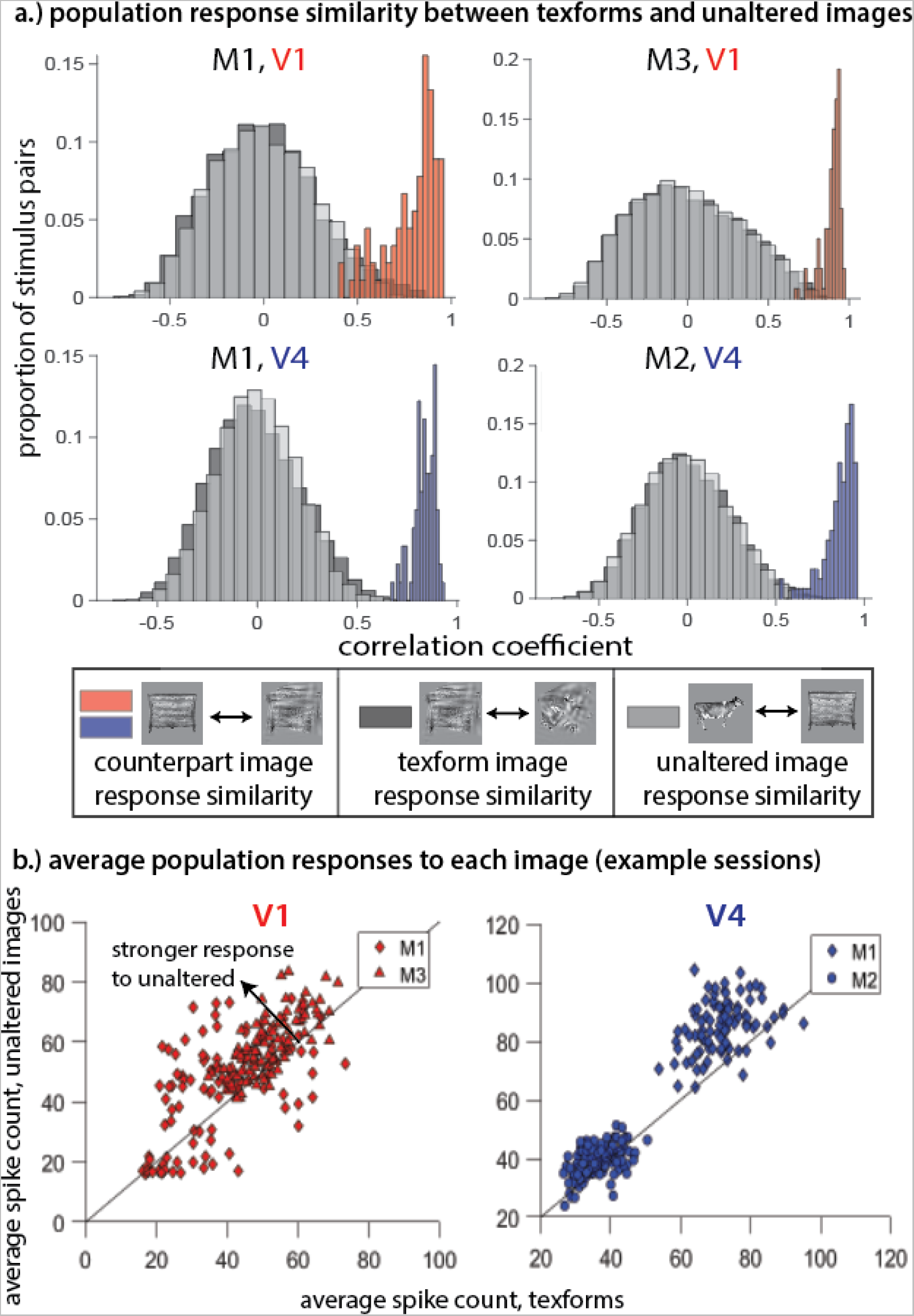
**(a) Example session population response similarity between texforms and unaltered images from example sessions.** Neuron-by-neuron correlations for each stimulus pair were computed as follows: for each stimulus and each neuron, the average spike count was computed across presentations of that stimulus. Red and blue histograms are the correlations between all units in a given area during presentations of an unaltered stimulus and its texform counterpart. Dark gray histograms are the correlations between units for all possible pairings of unaltered stimuli, and light gray histograms are the correlations for all possible pairings of unaltered stimuli with all texforms that are not its counterpart. **(b) Average population responses to each image.** Example session average stimulus evoked responses for each animal and brain area. A single point represents a stimulus pair (texform and unaltered counterpart). Average population responses were computed for each trial by taking the average spike count over all units in an area. Stimulus specific responses are the mean population responses across trials where a given stimulus is presented. For these example sessions, unaltered images elicited greater average population responses than texforms: paired t-tests: V1 (left): p=3.0 x 10^-4^ (M1) and p=1.4 x 10^-21^ (M3); p=3.2 x 10^-24^ (M1) and p=1.0 x 10^-7^ (M2).

Consistent with previous results using functional imaging in humans (Long et al., 2018), average population responses were slightly greater to the unaltered images than their associated texforms (most points in figure 3b are above the diagonal). Unaltered images elicited significantly (*p<0.05*) greater mean responses than texforms in nearly all sessions and animals (two sessions in M2, V4 did not reach significance). Despite this difference in average response to unaltered and texform images, there was a strong image-by-image correlation between the mean population responses to the unaltered and texform versions of the same image (correlation coefficient between points in Figure 3b, left and right; for V1, *r* = 0.56 for M1, *r* = 0.74 for M3; for V4, *r* = 0.39 for M1, *r* = 0.50 for M2).

### Animacy and real-world size are encoded similarly for unaltered images and texforms

The unaltered images of animals and objects have clear contours and finer-grained details, while texforms retain only the summary statistics of local feature information. Thus, while animacy and real-world size decoding was successful for both unaltered and texform stimuli in most conditions, this may be accomplished through sensitivity to different features encoded across the neural populations. Alternatively, it may be the case that the features supporting animacy and object size distinctions are similar between texforms and their unaltered counterparts.

To determine if the features supporting classification were similar for unaltered and texform stimuli, we trained our decoders for animacy (Figures 4a, b, left columns) or real-world size (Figures 4a, b, right columns) on unaltered images and tested them on texforms (Figure 4a) or vice versa (Figures 4b). We used a cross-validation procedure that required decoders to generalize to completely different images (i.e. we did not include unaltered or texform counterparts of images we were testing in the training sets). In both brain areas and in all monkeys, our ability to decode animacy and size remained above chance after this decoder swapping procedure (distributions of session performances lie above horizontal chance performance line, Figure 4a, b; t-tests, *p<0.05*). However, this swapping procedure caused a reduction in decoding performance compared to decoding texforms on a decoder trained on other texforms or unaltered images on a decoder trained on other unaltered images (most points below the diagonal in Figures 4a, b, c and d; t-tests, *p<0.05*). This combination of results suggests that at least some part of the signals related to animacy and real-world size arise from the responses of V1 and V4 neurons to low or mid-level image features that are preserved in the texforms and are diagnostic of category.

**Figure 4.**
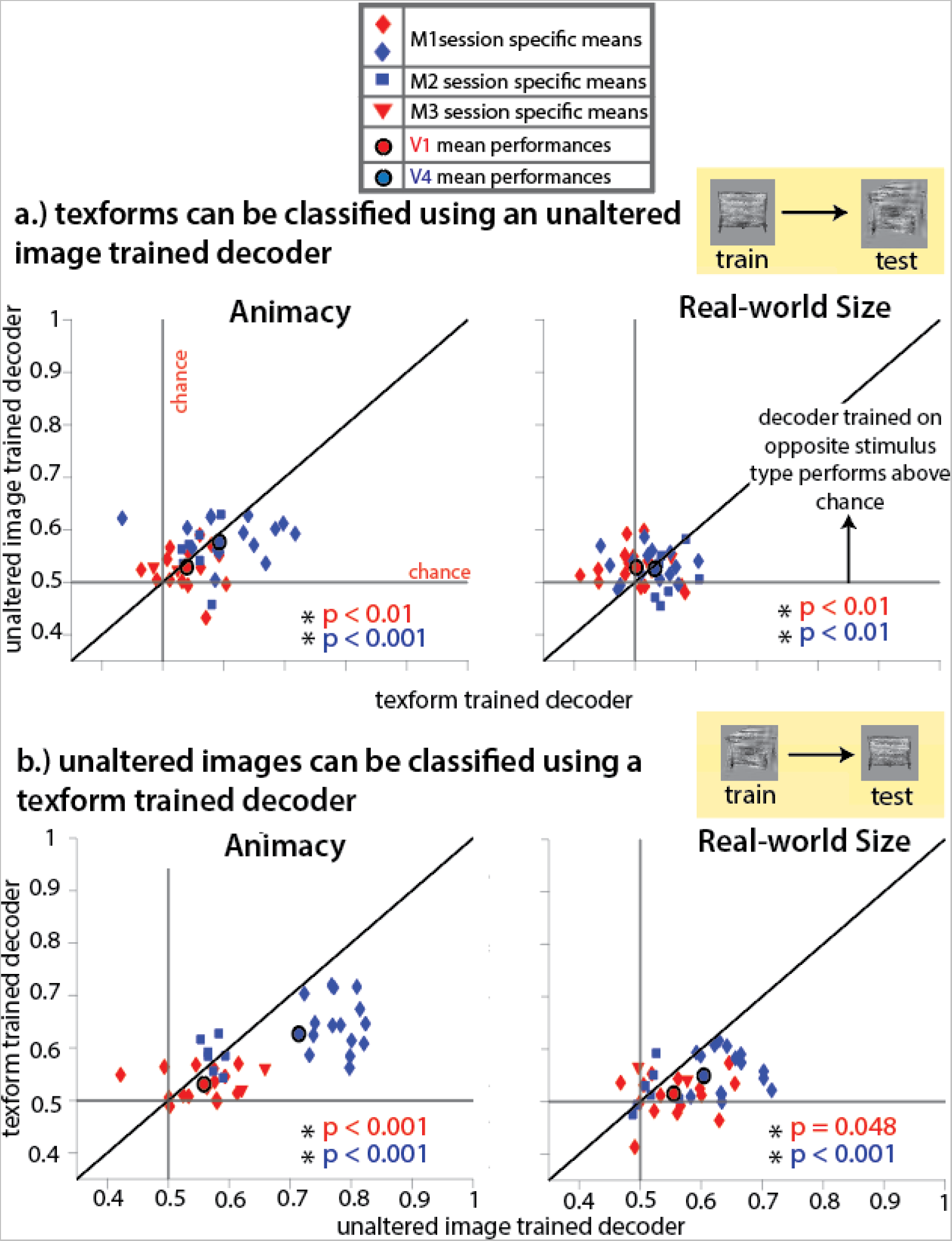
**(a) Texforms can be classified using an unaltered image trained decoder**. Performances of a linear decoder tasked with classifying the animacy (right) or real-world size (left) of texforms after being trained on trials corresponding to all other texforms (x axis) or unaltered images (y axis). Each animal specific symbol represents the mean decoding performance for one recording session. Each red and blue circle represents the mean decoding performance for a given brain area (V1 red, V4 blue), calculated by averaging over all sessions. Asterisks represent conditions where distributions of session decoding performances are significantly above chance when testing texform classification on unaltered image trained decoders. **(b) Unaltered images can be classified using a texform trained decoder.** Performances of a linear decoder tasked with classifying the animacy (right) or real-world size (left) of unaltered images after being trained on trials corresponding to all other unaltered images (x axis) or texforms (y axis). Each animal specific symbol represents the mean decoding performance for one recording session. Each red and blue circle represents the mean decoding performance for a given brain area (V1 red, V4 blue), calculated by averaging over all sessions. Asterisks represent conditions where distributions of session decoding performances are significantly above chance when testing unaltered image classification on texform trained decoders.

### Temporal analysis reveals no significant difference between transient and post-transient decoding performances for texforms or unaltered images

To further determine how the low- and mid-level visual features encoded by early and mid-level visual neurons contribute to animacy and real-world size categorization, we examined if and how decoding performances changed over the time course of the stimulus presentation period. A number of recent studies (Cauchoix et al., 2016; Wang et al., 2022) have sought to examine how well category information can be read out from the responses of mid-level visual neurons over time. One temporal prediction we sought to test posits that real-world size and animacy information is only decodable from the responses of mid-level visual neurons to texforms because of feedback from brain areas later in the visual stream (e.g. IT), reflecting further hierarchical processing (Grootswagers et al., 2019). Under this prediction, one would expect to see a gradual increase in the ability to decode animacy and real-world size of texforms as visual responses evolve, indicative of the restructuring of responses due to feedback from later processing stages.

To test if this prediction was accurate, we performed a temporal analysis where we compared the ability to decode animacy and real-world size using spike counts from either the transient or post-transient epochs of the response for both unaltered images and texforms (Figure 4b, c). The transient epoch was defined separately for each brain area, beginning at response latency (30 ms after stimulus onset for V1, and 50 ms for V4) and ending 100 ms later. Because a number of measures have proven sensitive to differences in firing rate (Churchland et al., 2010; Cohen & Kohn, 2011), we aimed to avoid biasing our results towards the transient decoder due to increased firing rates during the transient epoch compared to all other time points in the response. We opted to determine the length of the post-transient epoch separately for each session, animal, and brain area to ensure that the average number of spikes over the population was matched for transient and post-transient epochs (gray gradient, Figure 4a).

Although limiting spike counts to transient and post-transient windows did generally decrease decoding performances, there was no significant difference between the ability to decode animacy and real-world size information from V1 and V4 during these different time points for texforms or unaltered images (paired, two-sided Wilcoxon signed rank tests, p>0.05). This result implies that the initial (feedforward) responses of V1 and V4 neurons make significant contributions towards untangling representations of animacy and real-world size.

## Discussion

We demonstrated that groups of neurons in early and mid-level visual cortex encode information useful for categorizing objects according to their animacy and real-world size. Our ability to linearly decode animacy and real-world size information from monkey visual cortex was related to the ability of human observers to classify texforms by those same distinctions. These results indicate that signals that are directly useful for at least some higher-level distinctions are present in the early stages of visual processing. These signals might mediate the impressive speed and accuracy by which primates can categorize objects into ecologically important groups (Long et al., 2016, 2018; Yetter et al., 2021; Zachariou et al., 2018).

### A link between perception and linear decoding of neural population responses

Our finding that object categories can be, to an extent, linearly decoded even from groups of neurons early in the ventral stream suggests that animacy and real-world size are linearly separable, or partially untangled (DiCarlo & Cox, 2007), fairly early in processing. While animacy and real-world size might intuitively be assumed to be high-level, abstract concepts, our results suggest an alternative: low or mid-level visual features might 1) systematically vary across categories, 2) be encoded by early and mid-level ventral stream areas, and 3) be leveraged by the visual system to enable the ability to rapidly and accurately perceptually categorize objects.

Our results highlight a robust but not trivial link between linear decoding of neural population responses and perception. There is no a priori reason that the ability to decode the category of texforms from the responses of a few dozen V1 or V4 neurons in a monkey should be related to the average ability of humans to classify those texforms (Fig. 2b). The relationship between neural responses in early visual cortex and perception might have been profoundly nonlinear with a negligible linear component. The observation that the performance of a linear decoder of a few dozen monkey neurons is related to human perception suggests that linear decoding is a principle that is related to perception.

Further evidence for the value of low-level features for making what were thought to be primarily high-level distinctions comes from the converse of our finding: neurons even in late stages of the ventral visual stream are tuned for low-level features. For example, late stage ventral areas in humans and monkeys show tuning for a variety of low-level features including retinal position (DiCarlo & Maunsell, 2003), luminance (Baldassi et al., 2013), spatial frequency (Bonhoeffer & Grinvald, 1991), color (Conway & Tsao, 2009; Zeki, 1973), curvature (Yue et al., 2014, 2020), and the orientation of edges (D. S. P. Li & Bonner, 2021; Nasr et al., 2014; Nasr & Tootell, 2012). In human occipito-temporal cortex (OTC; a late stage ventral stream area), tuning for curvature, which is a mid-level feature most associated with selectivity in mid-level areas like V4 (Gallant et al., 1996; Pasupathy & Connor, 1999, 2001), accounted for about half of the variance in OTC responses to texforms (Long et al., 2018).

An interesting idea for future investigation is that some categories, perhaps those that are more important to the animal and/or differ more in low-level visual features, are linearly decodable (untangled) early in the visual system while others emerge only later. A recent study showed that without training, naive monkeys can discriminate features such as curvilinearity that are important for categories like animacy (Yetter et al., 2021), which they hypothesize is at least in part responsible for the remarkable speed and accuracy with which primates can categorize animacy.

### Looking forward: merging neuronal population recordings with human psychophysics can reveal fundamental properties of neural coding

Identifying the relationship between neural population responses and visually guided behaviors in behaving animals has led to many fundamental insights (Crapse & Basso, 2015; Cumming & Nienborg, 2016; Nienborg et al., 2012; Parker & Newsome, 1998; Ruff et al., 2018). One challenge of this approach is that the origin of those signals is difficult to interpret: in many cases, bottom-up, visual signals can be difficult to distinguish from signals of related to the animal’s internal state or other signals that are cognitive in origin (Chicharro et al., 2021; Cumming & Nienborg, 2016; Haefner et al., 2013; Nienborg & Cumming, 2010, 2009).

Comparing human psychophysics with neuronal population recordings in animals might be a way to distinguish those possibilities. Because the monkey and human subjects in our experiments cannot share variability in internal states, the link between recordings from monkey neurons and human psychophysics must mean that both reflect the image properties that guide the responses of visual neurons.

Our results highlight the value of comparing results using different experimental methodologies and different species to extract general principles about how the brain guides behavior. Doing so gives scientists the ability to leverage the unique strengths of each approach and system to illuminate the brain mechanisms that subserve a wide variety of perceptual and cognitive functions.

## Methods

### Animals

Our subjects were three adult male rhesus monkeys: M1, M2, M3 (*Macaca mulatta*, weighing 10, 10, and 11 kilograms, respectively). All animal procedures were approved by the Institutional Animal Care and Use Committees of the University of Pittsburgh and Carnegie Mellon University. Before any training or recording, animals received titanium head post implants. All monkeys had previously been used in tasks and had undergone prior electrophysiological recordings.

### All stimuli

Stimuli consisted of texforms and unaltered images. Before texforms were generated, unaltered stimuli were resized so that all images fit into the same size of (invisible) square on a gray background. This procedure ensured that real-world size categorization had to occur through the visual features that composed the image, not the size of the image on the screen. A total of 480 unique stimuli were used. Half of those stimuli were texforms, and the other half were the unaltered counterparts. Half of all stimuli (texforms and unaltered images) were taken from the stimulus set used in Long et al (2018), and the other half were taken from a stimulus set generated via the accelerated texform method (Deza et al., 2019). The stimulus set consisted of 60 images of big animals, 60 images of small animals, 60 images of big objects, and 60 images of small objects. For any given recording session, pairs of unaltered images and their texform counterparts were randomly selected for presentation to the animal, so that counts of stimuli in each category were roughly matched.

### Texform Stimuli

Texform stimuli were generated using a texture-synthesis algorithm via the following procedure: using the SHINE toolbox (Willenbockel et al., 2010), unaltered images were normalized for contrast and luminance (across the entire unaltered stimulus set). Each unaltered image was then placed in a larger gray display at a peripheral location so that it was encompassed by local spatial pooling windows generated by the algorithm. Thousands of first and second-order image statistics were then calculated within those local spatial pooling windows. The algorithm then iteratively coerced white noise to match those measured image statistics within the window. See <u>Long et al</u> (2018) and <u>texform generation code</u> resource page for an extensive explanation.

### Human Classification Scores

To determine if there is a relationship between the ability to decode animacy and size information from monkey visual neuron populations and humans classification performance, we chose to compare the neural results collected in this study with human behavioral results collected in previous studies (Chen, Deza, Long, & Konkle, unpublished data; Long et al., 2018 used similar methods, human studies conducted online via Amazon Mechanical Turk). In those previously conducted studies, texform stimuli were assigned classifiability scores based on humans’ ability to classify them correctly. All human psychophysical data was collected with approval from the Institutional Review Board at Harvard University and all subjects provided informed consent. While viewing texform stimuli, human participants responded yes or no to the questions below. In order to gauge animacy classification, one group of participants (N=16) were asked: “Here is a scrambled picture of something. Was the original thing an animal?”. Three other groups of participants (N=16 for each of the three groups) were asked: whether the texform was man made, whether it was small enough to be held with one or two hands, or whether it was large enough to support a human. Classification scores for each texform were then generated by subtracting the percent of incorrect classifications from the percent of correct classifications. For example, if a texform was generated from a picture of an animal, the score for animacy classification would be the percentage of responses indicating “yes, it’s an animal” subtracted from the percentage of responses indicating “yes, it’s a man made object”.

### Monkey Behavioral Task

Visual stimuli were displayed using a custom software (written in MATLAB using Psychophysics Toolbox (v.3), Brainard, 1997) on a CRT monitor (1,024x768 pixels; 120 Hz refresh rate) that was placed 54 cm away from each head fixed animal. Eye position was recorded using an infrared eye tracker (EyeLink 1000; SR Research). We recorded neuronal responses and the signal from a photodiode (used to align the neuronal responses to the stimulus presentation period) using Ripple hardware. Receptive field mapping occurred after array implantation, and once per week of recording. We did not observe considerable changes in receptive field location over the duration of the recording period. Stimuli were positioned on the screen so that their placement overlapped as many receptive fields as possible.

To initiate a trial, the animals were required to fixate on a small, central spot of light for 100-200 ms, and maintain that fixation while the stimulus (either an unaltered image or texform) was presented for a duration of 860 ms. Following stimulus offset, the fixation spot disappeared and a target appeared (target position sampled randomly from distribution of angles around the fixation point, but never overlapping the stimulus location). The animal received a liquid reward for making a saccade to the target (eye movement provided as a control for low-level adaptation). One stimulus was presented per trial. Little training was required (< 1 week, if any) to achieve adequate task performance for all animals.

Stimuli were presented to the animals in two possible sequences. The first sequence involved pairs of trials in which a unique stimulus would be presented back to back. There were four different conditions in this sequence, which depended on the ordering of the type (unaltered image=U or texform=T) of the stimuli in the pair. Therefore, each unique stimulus was presented under the following conditions: U/U, T/T, U/T, T/U. The second sequence type was in a four block design, with the first block being the presentation of 48 randomly ordered unique unaltered images and the second block being the previous block’s texform counterparts (presented in the same order as the unaltered stimuli). The third and fourth blocks used 48 different randomly ordered stimuli in the same manner, but presented the texform block before the unaltered stimuli block.

### Electrophysiological Recordings

Simultaneous extracellular recordings using chronically implanted 6x8 microelectrode arrays (Blackrock Microsystems, arrays placed in one hemisphere) were obtained in monkey M1 (visual areas V1 and V4, one array in each area) and monkey M2 (visual area V4, data from one of two arrays implanted analyzed in this study). The arrays were connected to a percutaneous pedestal which allowed for recording. Each electrode within the array was 1mm long, and the distance between adjacent electrodes was 400 μm. In monkey M3, we recorded from visual area V1 using linear, 32-channel moveable probes (Plexon) with an interelectrode spacing of 50 μm. Each area dataset (V1 or V4) contained data from two monkeys.

We defined analyzable recording sessions as those in which *at least* 60 unique stimuli with different identities (i.e. unaltered hawk/texform hawk, unaltered jukebox/texform jukebox, etc.) were presented to the animal. This criterion resulted in 15 recording sessions for M1, seven sessions for M2, and two sessions for M3. We recorded a combination of single neuron and multiunit clusters. We use the term “unit” to refer to either. Analyses comparing single and multiunits in our previous work (Cohen & Maunsell, 2009) as well as in another laboratory (Trautmann et al., 2019), did not find systematic differences between single neurons and multiunits for the types of population analyses presented here. For that reason, we analyzed multi-unit activity (one multiunit per channel). To determine which units to include from each recording session, we required that the transient response to the stimulus was at least 25% more than baseline activity (measured in the 100 ms post-fixation, pre-stimulus period). Units that did not reach this criterion were excluded from further analysis. The average number of included units per recording session for each monkey was: M1: 46 V1 units and 48 V4 units, M2: 35 V4 units, M3: 32 V1 units. In order to control for artifact noise that may be shared across multiple channels, we chose to exclude trials where the activity of more than 50% of units was greater than three standard deviations away from the mean activity for that unit.

### Statistical Analysis

We conducted most analyses using the spike count responses during the stimulus presentation period of each trial, after the expected response latency for each area (V1= 30ms, V4=50 ms). Only trials in which fixation was maintained for the entirety of the stimulus presentation period were included.

We compared the average population activity in each area for unaltered images and their texform counterparts to examine general differences in how each area responded to the different stimulus types. Average population response was quantified as the mean spike count across all units (in that area) for all presentations of that unique stimulus.

### Decoding Analyses

We used cross-validated linear decoding methods because we simply wanted to determine our ability to differentiate neuronal population responses between two categories. All decoders were determined separately for each animal, brain area, and recording session, and utilized spike counts (over the duration of the stimulus presentation period for general decoding performances, Figures 1, 2, 4; and over specific epochs for temporal analyses Fig. 5).

**Figure 5.**
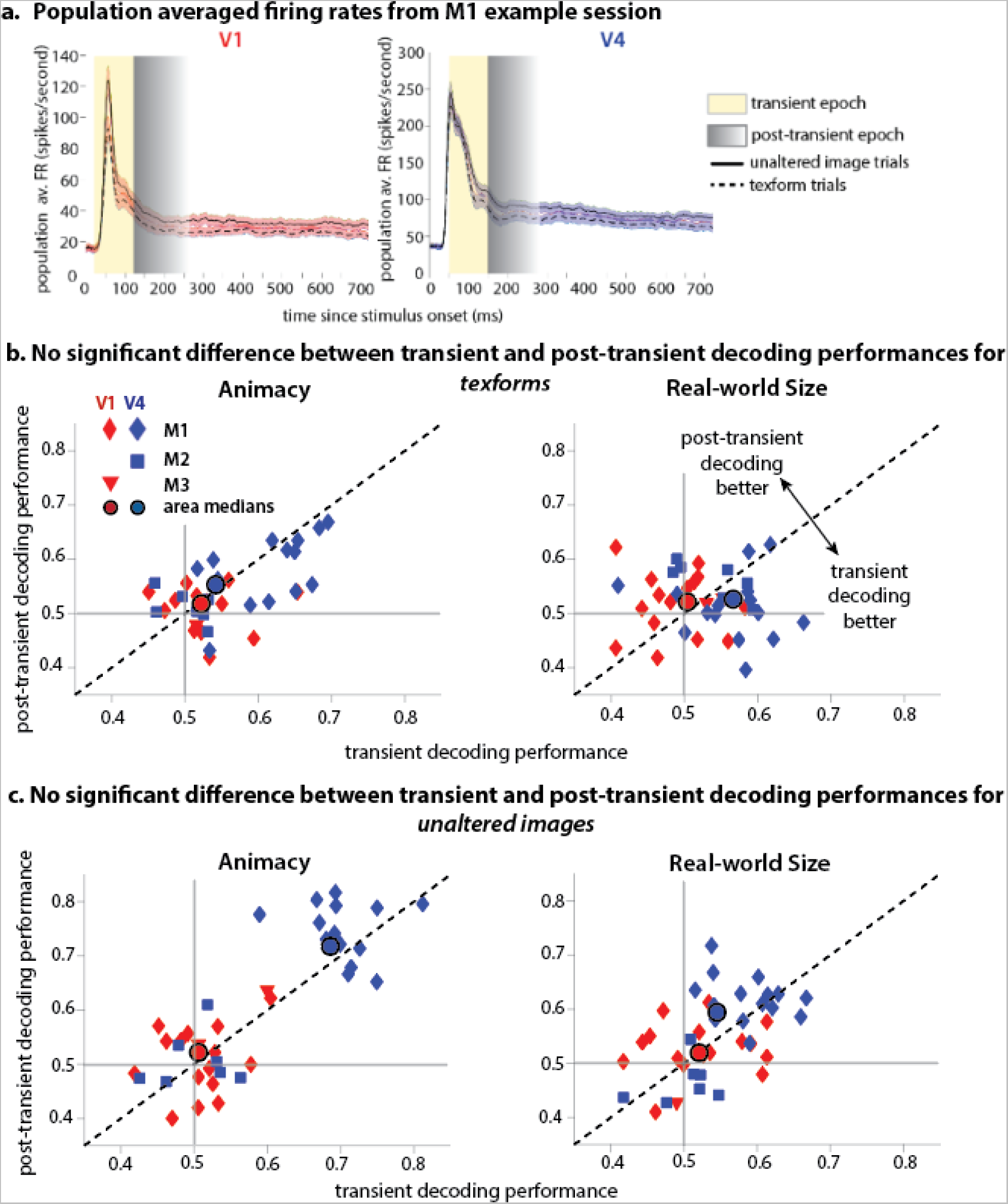
**(a). M1 example session population averaged firing rates** during the stimulus presentation period, for unaltered image trials (solid lines) and texform trials (dashed lines). Shaded regions represent standard error of the mean. The transient epoch (yellow) begins at response latency (V1=30 ms, V4=50 ms) and ends 100 ms later. The post-transient epoch (gray gradient) begins one millisecond after the end of the transient and concludes when the number of spikes (summed across the population) is greater than the average number of spikes across the population during the transient period. Post-transient epoch durations were calculated separately for unaltered images and texforms, and were determined on a session-by-session basis. **(b.) Decoding performances for unaltered images using transient and post-transient epoch spike counts.** Scatter plots show transient epoch decoding performances (x axis) and post-transient decoding performances (y axis) for animacy (left plot) and real-world size (right plot) for unaltered images. Each data point (blue and red animal specific symbols) represents the mean decoding performance (for transient and post-transient epochs) for each session, animal, and brain area. Circular data points represent the median transient and post-transient decoding performance (across animals and sessions) for each brain area. Paired, two-sided Wilcoxon ranked sign tests indicate no significant difference between decoding performances for transient and post-transient epochs for unaltered images (p>0.05). **(c.) Decoding performances for texforms using transient and post-transient epoch spike counts.** Scatter plots are formatted the same as those in (b.). Two-sided, paired Wilcoxon signed rank tests indicate no significant difference between decoding performances for transient and post-transient epochs for texforms (p>0.05).

#### Decoding Texform and Unaltered Image Size and Animacy

To assess the amount of information about size and animacy in our recorded neuronal populations, we trained and tested linear decoders separately on responses to texforms and unaltered stimuli. We used leave-two-out cross-validation in which we trained a decoder on responses to all but two images (one from each animacy or real-world size category), and determined the proportion of trials that were correctly categorized. This procedure prevents bias in the decoder when there are unequal numbers of trials in the two categories. We excluded all trials associated with the tested stimuli in order to ensure that the decoder was relying on category-specific feature information to classify stimuli, rather than stimulus-specific features. We repeated the stimulus sampling, decoder training, and testing procedure 2000 times to adequately assess the decoding performance for each session. We then quantified decoding performance as the proportion of correctly classified trials, averaged across all sessions for both animals (M1 and M3 for V1, M1 and M2 for V4). We assessed the significance of decoding performance for each brain area using a one-sided t-test to determine whether mean performance was significantly above chance performance of 0.5.

#### Cross-decoding Analyses

To determine whether size and animacy are separated in neuronal population space in similar ways for texforms and unaltered stimuli, we used a cross-decoding analysis. We calculated our ability to decode texforms or unaltered images with a decoder trained on the opposite stimulus type. As above, to ensure that the visual features specific to a stimulus did not artificially inflate the decoding performance, we excluded trials associated with the counterparts of the test stimuli from the training group.

#### Determining the relationship between human classification scores and monkey neural decoding

We used cross-validated linear regression to calculate the relationship between monkey neuronal population responses and the ability of humans to classify the animacy or real world size of texforms. For each texform stimulus in each recording session, we used linear regression to relate the responses of the recorded units to all other texform stimuli (averaged over all presentations of the same stimulus to construct a #units x #texform stimuli -1 matrix) to the human animacy and size classification scores (1 X # size or animacy scores - 1). We used those weights to predict the animacy or size classification score for the held out texform using the average population response to all presentations of the held out stimulus (Supplemental Figure 2, left). We used ROC analysis to assess how well an ideal observer could classify animacy or size using the predicted classification scores for each texform stimulus (Supplemental Figure 2, right). This process left us with four conditions: decoding size information from the human size score axis, decoding size information from the human animacy score axis, decoding animacy information from the human size score axis, and decoding animacy information from the human animacy score axis. To determine if decoding performance was significantly improved when utilizing the human score derived axis that was congruent to the type of decoding being conducted, we compared the congruent and incongruent decoding performance session distributions using one-sided, paired-sample t-tests.

Congruent condition decoding performances were also compared with a shuffled control. To generate null distributions of shuffled decoding performances, we randomly assigned human classification scores to texform stimuli (100 times per session), and repeated the congruent decoding condition from the analysis described above. We then found the average (over the 100 iterations) shuffled decoding performance for each session. To determine if congruent condition decoding performances were significantly better than shuffled decoding performances across sessions, we performed one-sided, paired-sample t-tests.

#### Temporal Decoding Analyses

We compared the ability to decode animacy and real-world size categories from transient and post-transient responses in V1 and V4 populations. The size of the temporal window we summed spike counts over for the post-transient epoch was determined separately for each session, brain area, and stimulus type (texform or unaltered images), in order to roughly match the number of spikes over the population in transient and post-transient conditions. First, the average (across texform or unaltered image trials) spike count (summed across units in a population) during the transient epoch was found. We constructed a matrix of average (across trials of a stimulus type) spike counts for each unit and one millisecond time bin, beginning with the time bin immediately following transient epoch end and concluding at the stimulus offset (#units x #time bins). We then found the cumulative sum of average spike counts (across the population and time) as we stepped through each time bin in this matrix, and found the time bin where the average number of spikes first surpassed the average number of spikes during the transient epoch. We took this time as the end point of the post-transient period. The average time of post-transient epoch conclusion for each brain area and stimulus type is displayed in Table 1, below.

**Table 1.** Mean post-transient epoch conclusions times for each brain area and stimulus type. End times are expressed in time since stimulus onset. Means and standard error of means (SEM) were calculated over all sessions (across animals) for a given brain area.

| Brain area,<br>stimulus type | V1, texforms | V1, unaltered<br>images | V4, texforms | V4, unaltered<br>images |
| --- | --- | --- | --- | --- |
| Mean<br>post-transient end<br>time | 254 ms | 251 ms | 268 ms | 255 ms |
| SEM (+/-) | 4.09 ms | 4.17 ms | 6.53 ms | 5.23 ms |

After finding the post-transient epochs for each recording session, we performed the temporal decoding analysis on matrices of spike counts (#units x #trials) tallied over either the transient or post-transient period. Utilizing the same leave-two-out cross-validated procedure described above, we calculated animacy and real-world size decoding performances for each session, stimulus type, and temporal epoch combination.

## Acknowledgements

This work was supported by the Simons Foundation (Simons Collaboration on the Global Brain award 542961SPI to M.R.C), the National Eye Institute of the National Institutes of Health (awards R01EY022930, R01EY034723, and RF1NS121913 to M.R.C). B.L. received support from the National Institutes of Health (K99 HD108386-01). T.K. received support from NSF CAREER BCS-1942438. We thank Arturo Deza for providing the stimuli. We thank Douglas Ruff for assistance in data collection. We thank Cheng Xue, Douglas Ruff, Amy Ni, and Ramanujan Srinath for helpful comments and feedback throughout this project. We thank Karen McKracken for technical assistance.

## Author Contributions

L.E.K and M.R.C designed the experiments with input from T.K. L.E.K conducted the experiments and performed all data analysis. Y.C. collected human behavioral data. L.E.K, Y.C., B.L., T.K., and M.R.C. wrote the paper.

## REFERENCES

Arcaro, M. J., & Livingstone, M. S. (2017). A hierarchical, retinotopic proto-organization of the primate visual system at birth. ELife, 6, e26196. https://doi.org/10.7554/eLife.26196

Baldassi, C., Alemi-Neissi, A., Pagan, M., DiCarlo, J. J., Zecchina, R., & Zoccolan, D. (2013). Shape Similarity, Better than Semantic Membership, Accounts for the Structure of Visual Object Representations in a Population of Monkey Inferotemporal Neurons. PLoS Computational Biology, 9(8), e1003167. https://doi.org/10.1371/journal.pcbi.1003167

Biederman, I. (1987). Recognition-by-components: A theory of human image understanding. Psychological Review, 94(2), 115–147. https://doi.org/10.1037/0033-295X.94.2.115

Bonhoeffer, T., & Grinvald, A. (1991). Iso-orientation domains in cat visual cortex are arranged in pinwheel-like patterns. Nature, 353(6343), 429–431. https://doi.org/10.1038/353429a0

Brainard, D. H. (1997). The Psychophysics Toolbox. Spatial Vision, 10(4), 433–436. https://doi.org/10.1163/156856897X00357

Cauchoix, M., Crouzet, S. M., Fize, D., & Serre, T. (2016). Fast ventral stream neural activity enables rapid visual categorization. NeuroImage, 125, 280–290. https://doi.org/10.1016/j.neuroimage.2015.10.012

Chicharro, D., Panzeri, S., & Haefner, R. M. (2021). Stimulus-dependent relationships between behavioral choice and sensory neural responses. ELife, 10, e54858. https://doi.org/10.7554/eLife.54858

Churchland, M. M., Yu, B. M., Cunningham, J. P., Sugrue, L. P., Cohen, M. R., Corrado, G. S., Newsome, W. T., Clark, A. M., Hosseini, P., Scott, B. B., Bradley, D. C., Smith, M. A., Kohn, A., Movshon, J. A., Armstrong, K. M., Moore, T., Chang, S. W., Snyder, L. H., Lisberger, S. G., … Shenoy, K. V. (2010). Stimulus onset quenches neural variability: A widespread cortical phenomenon. Nature Neuroscience, 13(3), 369–378. https://doi.org/10.1038/nn.2501

Cohen, M. R., & Kohn, A. (2011). Measuring and interpreting neuronal correlations. Nature Neuroscience, 14(7), 811–819. https://doi.org/10.1038/nn.2842

Cohen, M. R., & Maunsell, J. H. R. (2009). Attention improves performance primarily by reducing interneuronal correlations. Nature Neuroscience, 12(12), 1594–1600. https://doi.org/10.1038/nn.2439

Conway, B. R., & Tsao, D. Y. (2009). Color-tuned neurons are spatially clustered according to color preference within alert macaque posterior inferior temporal cortex. Proceedings of the National Academy of Sciences, 106(42), 18034–18039. https://doi.org/10.1073/pnas.0810943106

Crapse, T. B., & Basso, M. A. (2015). Insights into decision making using choice probability. Journal of Neurophysiology, 114(6), 3039–3049. https://doi.org/10.1152/jn.00335.2015

Cumming, B. G., & Nienborg, H. (2016). Feedforward and feedback sources of choice probability in neural population responses. Current Opinion in Neurobiology, 37, 126–132. https://doi.org/10.1016/j.conb.2016.01.009

Deza, A., Chen, Y.-C., Long, B., & Konkle, T. (2019). Accelerated Texforms: Alternative Methods for Generating Unrecognizable Object Images with Preserved Mid-Level Features. 2019 Conference on Cognitive Computational Neuroscience. 2019 Conference on Cognitive Computational Neuroscience, Berlin, Germany. https://doi.org/10.32470/CCN.2019.1412-0

DiCarlo, J. J., & Cox, D. D. (2007). Untangling invariant object recognition. Trends in Cognitive Sciences, 11(8), 333–341. https://doi.org/10.1016/j.tics.2007.06.010

DiCarlo, J. J., & Maunsell, J. H. R. (2003). Anterior Inferotemporal Neurons of Monkeys Engaged in Object Recognition Can be Highly Sensitive to Object Retinal Position. Journal of Neurophysiology, 89(6), 3264–3278. https://doi.org/10.1152/jn.00358.2002

Felleman, D. J., & Van Essen, D. C. (1991). Distributed hierarchical processing in the primate cerebral cortex. *Cerebral Cortex (New York*, N.Y.: 1991), *1*(1), 1–47. https://doi.org/10.1093/cercor/1.1.1-a

Freeman, J., & Simoncelli, E. P. (2011). Metamers of the ventral stream. Nature Neuroscience, 14(9), 1195–1201. https://doi.org/10.1038/nn.2889

Gallant, J. L., Connor, C. E., Rakshit, S., Lewis, J. W., & Van Essen, D. C. (1996). Neural responses to polar, hyperbolic, and Cartesian gratings in area V4 of the macaque monkey. Journal of Neurophysiology, 76(4), 2718–2739. https://doi.org/10.1152/jn.1996.76.4.2718

Grill-Spector, K., & Weiner, K. S. (2014). The functional architecture of the ventral temporal cortex and its role in categorization. Nature Reviews Neuroscience, 15(8), 536–548. https://doi.org/10.1038/nrn3747

Grootswagers, T., Robinson, A. K., Shatek, S. M., & Carlson, T. A. (2019). Untangling featural and conceptual object representations. NeuroImage, 202, 116083. https://doi.org/10.1016/j.neuroimage.2019.116083

Haefner, R. M., Gerwinn, S., Macke, J. H., & Bethge, M. (2013). Inferring decoding strategies from choice probabilities in the presence of correlated variability. Nature Neuroscience,16(2), 235–242. https://doi.org/10.1038/nn.3309

Hasson, U., Levy, I., Behrmann, M., Hendler, T., & Malach, R. (2002). Eccentricity Bias as an Organizing Principle for Human High-Order Object Areas. Neuron, 34(3), 479–490. https://doi.org/10.1016/S0896-6273(02)00662-1

Khaligh-Razavi, S.-M., & Kriegeskorte, N. (2014). Deep Supervised, but Not Unsupervised, Models May Explain IT Cortical Representation. PLoS Computational Biology, 10(11), e1003915. https://doi.org/10.1371/journal.pcbi.1003915

Kravitz, D. J., Saleem, K. S., Baker, C. I., Ungerleider, L. G., & Mishkin, M. (2013). The ventral visual pathway: An expanded neural framework for the processing of object quality. Trends in Cognitive Sciences, 17(1), 26–49. https://doi.org/10.1016/j.tics.2012.10.011

Krizhevsky, A., Sutskever, I., & Hinton, G. E. (2012). ImageNet Classification with Deep Convolutional Neural Networks. In F. Pereira, C. J. Burges, L. Bottou, & K. Q. Weinberger (Eds.), Advances in Neural Information Processing Systems (Vol. 25). Curran Associates, Inc. https://proceedings.neurips.cc/paper/2012/file/c399862d3b9d6b76c8436e924a68c45b-Paper.pdf

Li, D. S. P., & Bonner, M. F. (2021). *Emergent selectivity for scenes, object properties, and contour statistics in feedforward models of scene-preferring cortex* [Preprint].Neuroscience. https://doi.org/10.1101/2021.09.24.461733

Li, S. P., & Bonner, M. (2021). Deep neural network models of visual cortex reveal curvature and real-world size as organizing principles of mid-level representation. Journal of Vision, 21(9), 2751. https://doi.org/10.1167/jov.21.9.2751

Lieber, J. D., Lee, G. M., Majaj, N. J., & Movshon, J. A. (2023). Sensitivity to naturalistic texture relies primarily on high spatial frequencies. Journal of Vision, 23(2), 4. https://doi.org/10.1167/jov.23.2.4

Long, B., Konkle, T., Cohen, M. A., & Alvarez, G. A. (2016). Mid-level perceptual features distinguish objects of different real-world sizes. Journal of Experimental Psychology: General, 145(1), 95–109. https://doi.org/10.1037/xge0000130

Long, B., Störmer, V. S., & Alvarez, G. A. (2017). Mid-level perceptual features contain early cues to animacy. Journal of Vision, 17(6), 20. https://doi.org/10.1167/17.6.20

Long, B., Yu, C.-P., & Konkle, T. (2018). Mid-level visual features underlie the high-level categorical organization of the ventral stream. Proceedings of the National Academy of Sciences, 115(38). https://doi.org/10.1073/pnas.1719616115

Mishkin, M., Ungerleider, L. G., & Macko, K. A. (1983). Object vision and spatial vision: Two cortical pathways. Trends in Neurosciences, 6, 414–417. https://doi.org/10.1016/0166-2236(83)90190-X

Nasr, S., Echavarria, C. E., & Tootell, R. B. H. (2014). Thinking Outside the Box: Rectilinear Shapes Selectively Activate Scene-Selective Cortex. Journal of Neuroscience, 34(20), 6721–6735. https://doi.org/10.1523/JNEUROSCI.4802-13.2014

Nasr, S., & Tootell, R. B. H. (2012). A Cardinal Orientation Bias in Scene-Selective Visual Cortex. Journal of Neuroscience, 32(43), 14921–14926. https://doi.org/10.1523/JNEUROSCI.2036-12.2012

Nienborg, H., & Cumming, B. (2010). Correlations between the activity of sensory neurons and behavior: How much do they tell us about a neuron’s causality? Current Opinion in Neurobiology, 20(3), 376–381. https://doi.org/10.1016/j.conb.2010.05.002

Nienborg, H., & Cumming, B. G. (2009). Decision-related activity in sensory neurons reflects more than a neuron’s causal effect. Nature, 459(7243), 89–92. https://doi.org/10.1038/nature07821

Nienborg, H., R. Cohen, M., & Cumming, B. G. (2012). Decision-Related Activity in Sensory Neurons: Correlations Among Neurons and with Behavior. Annual Review of Neuroscience, 35(1), 463–483. https://doi.org/10.1146/annurev-neuro-062111-150403

Oleskiw, T. D., Lieber, J. D., Movshon, J. A., & Simoncelli, E. P. (2020). Testing a two-stage model of stimulus selectivity in macaque V2. Journal of Vision, 20(11), 1540. https://doi.org/10.1167/jov.20.11.1540

Parker, A. J., & Newsome, W. T. (1998). Sense and the single neuron: Probing the physiology of perception. Annual Review of Neuroscience, 21, 227–277. https://doi.org/10.1146/annurev.neuro.21.1.227

Pasupathy, A., & Connor, C. E. (1999). Responses to Contour Features in Macaque Area V4. Journal of Neurophysiology, 82(5), 2490–2502. https://doi.org/10.1152/jn.1999.82.5.2490

Pasupathy, A., & Connor, C. E. (2001). Shape Representation in Area V4: Position-Specific Tuning for Boundary Conformation. Journal of Neurophysiology, 86(5), 2505–2519. https://doi.org/10.1152/jn.2001.86.5.2505

Riesenhuber, M., & Poggio, T. (1999). Hierarchical models of object recognition in cortex. Nature Neuroscience, 2(11), 1019–1025. https://doi.org/10.1038/14819

Ruff, D. A., Ni, A. M., & Cohen, M. R. (2018). Cognition as a Window into Neuronal Population Space. Annual Review of Neuroscience, 41(1), 77–97. https://doi.org/10.1146/annurev-neuro-080317-061936

Trautmann, E. M., Stavisky, S. D., Lahiri, S., Ames, K. C., Kaufman, M. T., O’Shea, D. J., Vyas, S., Sun, X., Ryu, S. I., Ganguli, S., & Shenoy, K. V. (2019). Accurate Estimation of Neural Population Dynamics without Spike Sorting. Neuron, 103(2), 292–308.e4. https://doi.org/10.1016/j.neuron.2019.05.003

Ungerleider, L. G., & Bell, A. H. (2011). Uncovering the visual “alphabet”: Advances in our understanding of object perception. Vision Research, 51(7), 782–799. https://doi.org/10.1016/j.visres.2010.10.002

Wang, R., Janini, D., & Konkle, T. (2022). Mid-level Feature Differences Support Early Animacy and Object Size Distinctions: Evidence from Electroencephalography Decoding. Journal of Cognitive Neuroscience, 34(9), 1670–1680. https://doi.org/10.1162/jocn_a_01883

Willenbockel, V., Sadr, J., Fiset, D., Horne, G. O., Gosselin, F., & Tanaka, J. W. (2010). Controlling low-level image properties: The SHINE toolbox. Behavior Research Methods, 42(3), 671–684. https://doi.org/10.3758/BRM.42.3.671

Yamins, D. L. K., & DiCarlo, J. J. (2016). Using goal-driven deep learning models to understand sensory cortex. Nature Neuroscience, 19(3), 356–365. https://doi.org/10.1038/nn.4244

Yamins, D. L. K., Hong, H., Cadieu, C. F., Solomon, E. A., Seibert, D., & DiCarlo, J. J. (2014). Performance-optimized hierarchical models predict neural responses in higher visual cortex. Proceedings of the National Academy of Sciences, 111(23), 8619–8624. https://doi.org/10.1073/pnas.1403112111

Yetter, M., Robert, S., Mammarella, G., Richmond, B., Eldridge, M. A. G., Ungerleider, L. G., & Yue, X. (2021). Curvilinear features are important for animate/inanimate categorization in macaques. Journal of Vision, 21(4), 3. https://doi.org/10.1167/jov.21.4.3

Yue, X., Pourladian, I. S., Tootell, R. B. H., & Ungerleider, L. G. (2014). Curvature-processing network in macaque visual cortex. Proceedings of the National Academy of Sciences,111(33). https://doi.org/10.1073/pnas.1412616111

Yue, X., Robert, S., & Ungerleider, L. G. (2020). Curvature processing in human visual cortical areas. NeuroImage, 222, 117295. https://doi.org/10.1016/j.neuroimage.2020.117295

Zachariou, V., Del Giacco, A. C., Ungerleider, L. G., & Yue, X. (2018). Bottom-up processing of curvilinear visual features is sufficient for animate/inanimate object categorization.Journal of Vision, 18(12), 3. https://doi.org/10.1167/18.12.3

Zeki, S. M. (1973). Colour coding in rhesus monkey prestriate cortex. Brain Research, 53(2), 422–427. https://doi.org/10.1016/0006-8993(73)90227-8

