## Supplemental Figures for "Contributions of early and mid-level visual cortex to high-level object categorization"

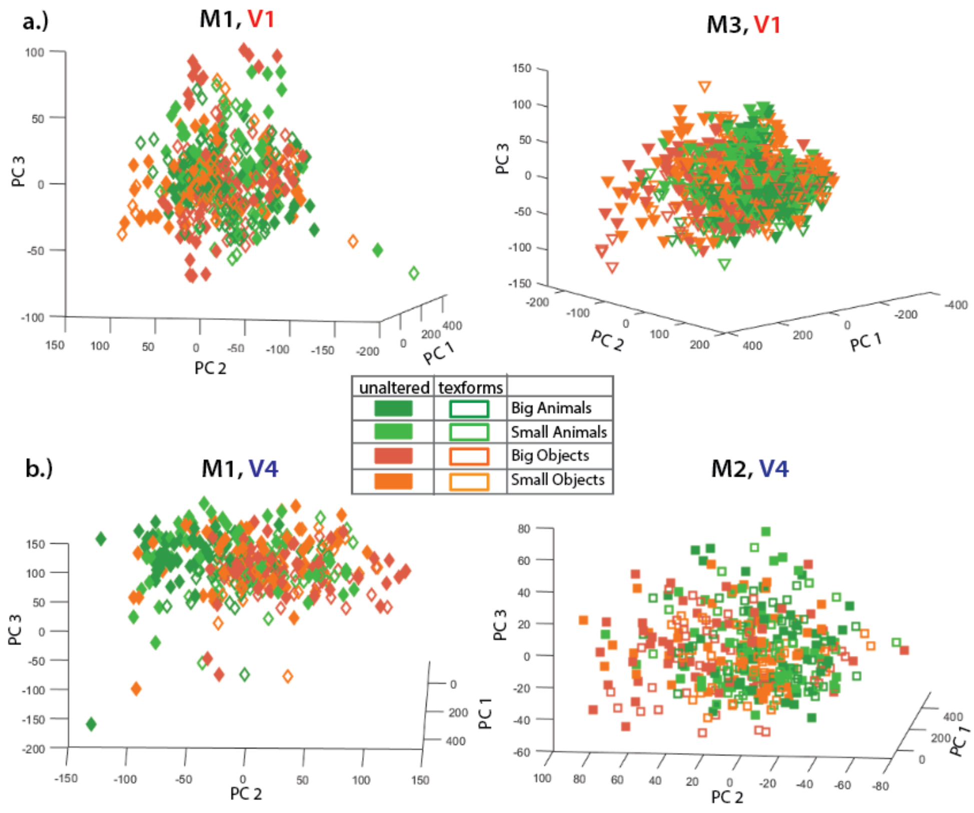


***Supplemental Figure 1. (a) Visualization of category separation in V1 PC space.*** *Each point represents the principal component score of a trial in an example session colored according to its category identity. Axes were rotated to best visualize the separation between categories.* ***(b) Visualization of category separation in V4 PC space.*** *Each point represents the principal component score of a trial in an example session colored according to its category identity. Axes were rotated to best visualize the separation between categories.*

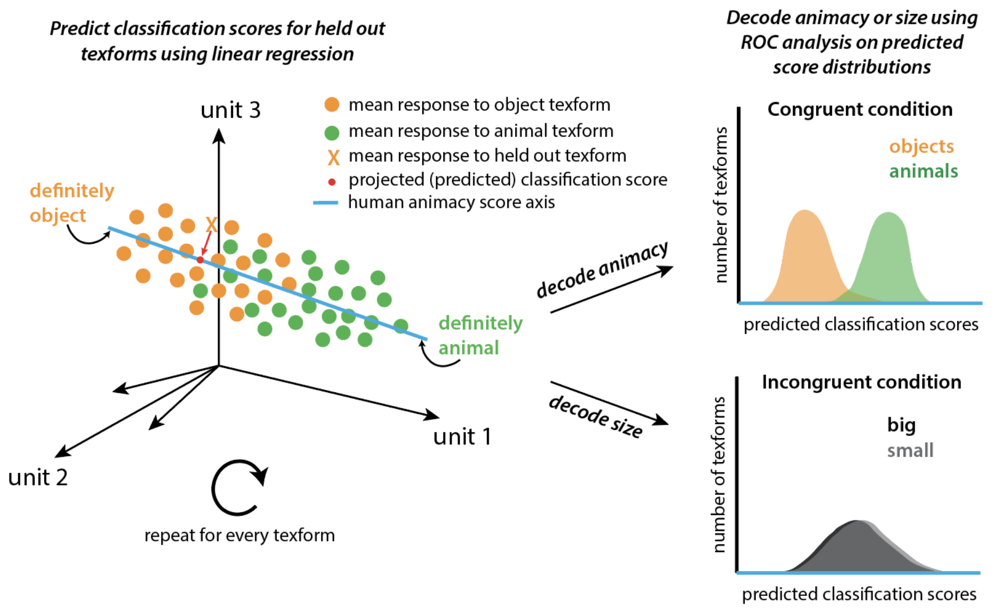


***Supplemental Figure 2. Schematic illustrating relationship between human classification scores and monkey neural decoding.*** *The figure depicts analytical methods for predicting animacy classification scores for all texforms presented in a given session (left), as well as the separation between distributions of those predicted scores when attempting to decode animacy (congruent human axis condition) or real-world size (incongruent human axis condition) information (right).*
